# Integrating structured and unstructured citizen science data to improve wildlife population monitoring

**DOI:** 10.1101/2021.03.03.431294

**Authors:** Philipp H. Boersch-Supan, Robert A. Robinson

**Affiliations:** British Trust for Ornithology, Thetford, United Kingdom; Department of Geography and Emerging Pathogens Institute, University of Florida, Gainesville, FL, United States

**Keywords:** biodiversity monitoring, breeding bird survey, Citizen science, population trend, data integration

## Abstract

Accurate and robust population trend assessments are key to successful biodiversity conservation. Citizen science surveys have provided good evidence of biodiversity declines whilst engaging people with them. Citizen scientists are also collecting opportunistic biodiversity records at unprecedented scales, vastly outnumbering records gathered through structured surveys. Opportunistic records exhibit spatio-temporal biases and heterogeneity in observer effort and skill, but their quantity offers a rich source of information. Data integration, the combination of multiple information sources in a common analytical framework, can potentially improve inferences about populations compared to analysing either in isolation. We combine count data from a structured citizen science survey and detection-nondetection data from an opportunistic citizen science programme. Population trends were modelled using dynamic N-mixture models to integrate both data sources. We applied this approach to two different inferential challenges arising from sparse data: (i) the estimation of population trends for an area smaller than a structured survey stratum, and (ii) the estimation of national population trends for a rare but widespread species. In both cases, data integration yielded population trajectories similar to those estimated from structured survey data alone but had higher precision when the density of opportunistic records was high. In some cases this allowed inferences about population trends where indices derived from single data sources were too uncertain to assess change. However, there were differences in the trend magnitude between the integrated and the standard survey model.

We show that data integration of large-scale structured and unstructured data is feasible and offers potential to improve national and regional wildlife trend estimates, although a need to independently validate trends remains. Smaller gains are achieved in areas where uptake of opportunistic recording is low. The integration of opportunistic records from volunteer-selected locations alone may therefore not adequately address monitoring gaps for management and policy applications. To achieve the latter, scheme organisers should consider providing incentives for achieving representative coverage of target areas in both structured and unstructured recording schemes.

## 1 Introduction

The ability to accurately and robustly quantify species’ population trajectories is key to successful biodiversity conservation. Monitoring of changes in a species’ population size or range is essential to assess threat status; to act as an early-warning signal of population declines; for conservation resource prioritization; and to assess the efficacy of environmental policies (Lawton 1993; Johnston *et al*. 2015). Yet, most wildlife populations cannot be completely enumerated, or even robustly surveyed, because resources for monitoring are finite. Large geographic biases exist in monitoring effort even for well sampled taxa like birds (Meyer *et al*. 2015, 2016; Amano, Lamming & Sutherland 2016). This affects our knowledge of species distributions, as well as our understanding of the processes underlying population dynamics because potential drivers, such as climate or land-use change, can differ between surveyed and unsurveyed regions (Pearce-Higgins *et al*. 2015).

National scale biodiversity monitoring schemes such as those that make up the Pan-European Common Bird Monitoring Scheme (van Strien, Pannekoek & Gibbons 2001; Birdlife International 2004) or the European Butterfly Monitoring Scheme (van Swaay *et al*. 2008, 2019) are designed to provide coverage of a broad range of common species, allowing the derivation of indicators of the state of nature while making the most of finite resources (Burns *et al*. 2018; Hayhow *et al*. 2019). Such high-level efforts have become closely intertwined with high-level (i.e. national and supra-national) conservation legislation and policy. However, the implementation of conservation policy on a legislative and executive level is increasingly devolved within nations. For example, in the UK conservation is now devolved to sub-national governments (NUTS 1 level) and their executive agencies, resulting in legislation and implementation approaches, including e.g. red list assessments, that are specific to England, Scotland, Wales, and Northern Ireland (Bainbridge 2014; Kirsop-Taylor 2019); similarly, in Germany federal conservation legislation provides an overarching legal framework but delegates implementation to the states, and the executive agencies implementing state laws may be devolved further to government regions (NUTS 2 level) or districts (NUTS 3 level) (Rose-Ackerman 1994). There is also a shift from treating conservation and management as jurisdictional issues towards more holistic approaches focussed on the maintenance of healthy ecosystems and ecosystem services at the appropriate spatial scales (Kirsop-Taylor 2019). Apart from individually designated large protected areas (e.g. national parks, protected landscapes), such approaches are promoted e.g. through the European Landscape Convention and within programmes of the EU common agricultural policy (Lomba *et al*. 2014). Again, the implementation varies among signatory states. Within the UK such natural subdivisions are reflected, for example, in the National Character Areas in England (Natural England 2014) or the Area Statements in Wales (Welsh Government 2017), both of which are based on a combination of landscape features, bio- and geodiversity, and socio-economic activity. The plethora of spatial units that arise from jurisdictional devolution and landscape-centric approaches, creates an increasing desire to repurpose data from national biodiversity monitoring schemes to provide information at smaller spatial scales, not addressed by national trends and indicators.

Many national biodiversity monitoring schemes are based on long-term structured surveys, which use predetermined monitoring sites and standardized survey methods. Structured surveys provide robust estimates of population trends but require large and long-term commitments by institutions – and where conducted as citizen science schemes, volunteers – and can be challenging to organize and coordinate (Schmeller *et al*. 2009). Instead, projects which rely on opportunistic records by interested members of the public may be a more effective means to increase the spatio-temporal coverage of distribution and abundance data (Dickinson, Zuckerberg & Bonter 2010; Isaac & Pocock 2015). Although such projects may have primary goals other than monitoring, e.g. raising awareness about focal taxa or facilitating personal record keeping for naturalists, there is increasing interest in using such schemes to fill knowledge gaps in regions that are poorly or not at all covered by structured surveys, and as a basis to obtain indices of population trajectories that meaningfully capture true wildlife population trends (Kéry *et al*. 2010; Isaac *et al*. 2014; Horns, Adler & Şekercioğlu 2018). Trend modelling based on such data is challenging because of known biases in site selection, visit timing, survey effort, and/or surveyor skill (Isaac & Pocock 2015; Johnston *et al*. 2018, 2020). Thus, there is usually a trade-off between collecting a small amount of higher ‘quality’ data conforming to a defined common structure or a larger amount of relatively heterogeneous (i.e. lower ‘quality’) data (Gardiner *et al*. 2012; Bayraktarov *et al*. 2018).

The consequences of this trade-off are a topic of active research (Aceves-Bueno *et al*. 2017; Bayraktarov *et al*. 2018; Kelling *et al*. 2018; Specht & Lewandowski 2018; Boersch-Supan, Trask & Baillie 2019; Johnston *et al*. 2020; Robinson *et al*. 2020), and there is a growing set of modelling approaches to address the challenges of unstructured datasets using auxiliary structured biodiversity data and/or observation models that account for preferential sampling, usually at the cost of increased model complexity and computational demands (van Strien, van Swaay & Termaat 2013; Fithian *et al*. 2015; Robinson, Ruiz-Gutierrez & Fink 2018; Isaac *et al*. 2019; Johnston *et al*. 2019, 2020).

Other recent work has investigated whether relatively simple models are sufficient to extract population trend information from less structured data (Roberts, Donald & Green 2007; Roy *et al*. 2012; Walker & Taylor 2017; Boersch-Supan *et al*. 2019). These simpler approaches generally rely on the assumption that the information gain from a larger quantity of records outpaces potential biases from opportunistic sampling.

However, integrating these two data sources may help overcome some of these issues, by combining the structure of survey data with the improved coverage of less structured schemes. This has the potential to improve the precision of model parameters and the resulting inferences (Fithian *et al*. 2015; Isaac *et al*. 2019), perhaps especially for species that are poorly covered by structured monitoring programmes, as well as offering a route to gain a more mechanistic understanding of the drivers of those population dynamics.

In the UK, biological recording by volunteers provides information on the occurrence or abundance of over 10000 taxa, although records are sparse for the vast majority of taxa (Roy *et al*. 2014; Hayhow *et al*. 2019; Outhwaite *et al*. 2020). As part of these efforts, comprehensive structured bird monitoring is undertaken through the Breeding Bird Survey (BBS; Figure 1) which provides population trends for about 120 common and widespread bird species (Greenwood *et al*. 1995; Freeman *et al*. 2007; Harris *et al*. 2018), but knowledge gaps remain for rare and cryptic species (approximately 220 species are regular breeders (Robinson 2010)).

**Figure 1:**
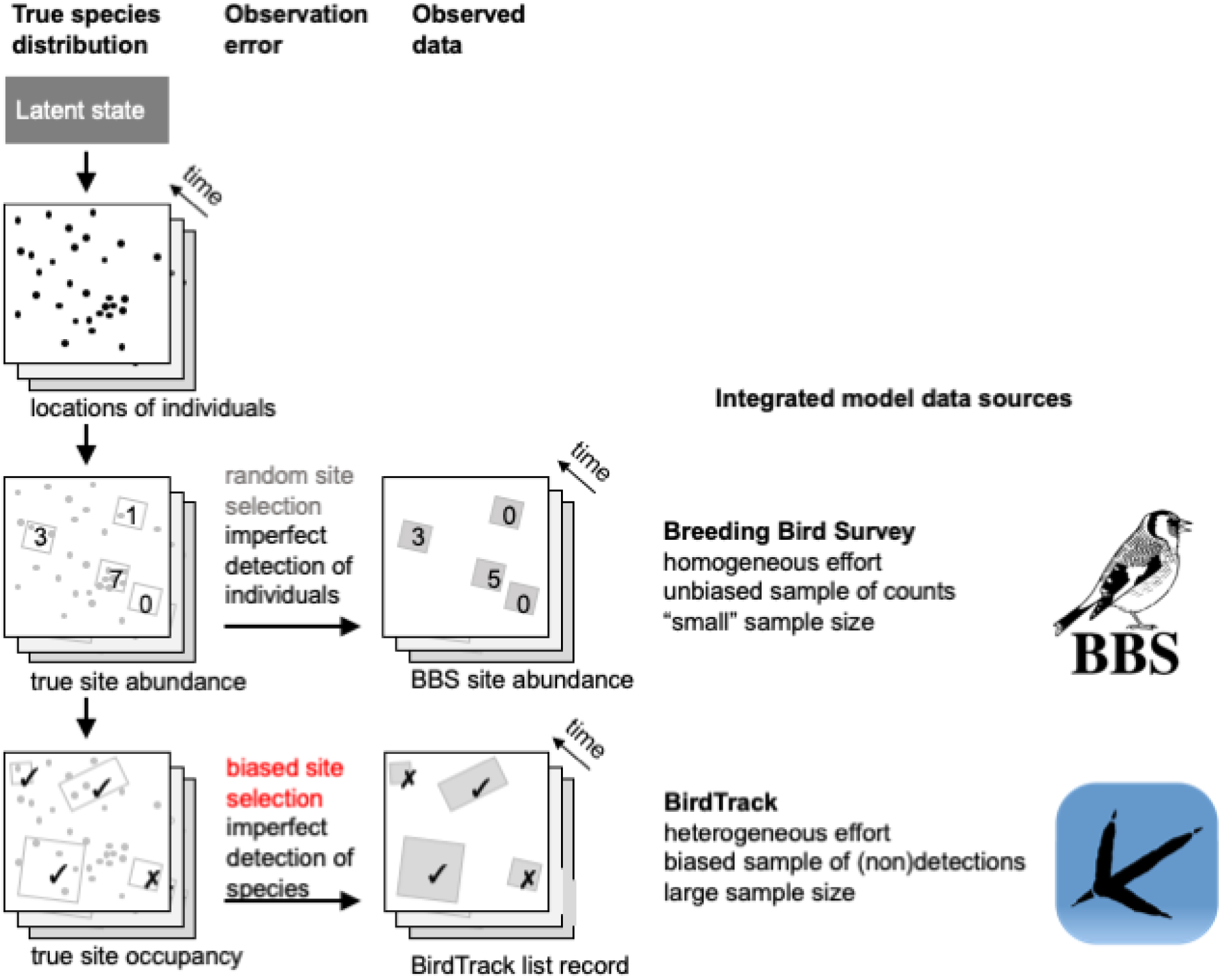
BBS surveys and BirdTrack lists are both observations of the true spatio-temporal distribution of birds. Observations from each scheme differ in their information quality and quantity. BBS counts are collected with known effort and spatially unbiased, but comparably sparse. BirdTrack lists are more numerous, but come from non-random locations and effort is heterogeneous. Figure adapted from Isaac et al. (2019).

Opportunistic citizen science recording schemes such as BirdTrack (Figure 1; www.birdtrack.net; Baillie *et al*. (2006); Newson *et al*. (2016)) provide greater coverage in space and time, but lack the structured protocols and formal sampling design. A recent comparison of these two datasets showed that national-scale annual reporting rate trends in BirdTrack were broadly consistent with BBS abundance trends for common species, and those exhibiting marked population changes (Boersch-Supan *et al*. 2019). However, the magnitude of reporting rate– abundance relationships were inconsistent across species, and agreement in trends for rarer species could not be ascertained, in part because of high uncertainty about population change in trends from either dataset.

In this study, we leverage the spatio-temporal overlap of two citizen science schemes to investigate the utility of joint analyses of structured and opportunistic datasets to derive population trends for uncommon breeding birds at regional and local scales. This has the potential to address a gap in currently available monitoring products with high relevance for landscape management. In particular we evaluate integrated trend models for two different inferential challenges: (i) the estimation of population trends for areas smaller than a BBS stratum to improve small-area inferences about population trends for local managers and decision makers, and (ii) the estimation of national population trends for a rare but widespread species to assess the utility of data integration when a species is not widespread enough to fulfil minimum sample size criteria for the structured survey model.

## 2 Materials and Methods

### 2.1 Data sources

We employed structured survey data from the Breeding Bird Survey (BBS) (Gregory, Baillie & Bashford 2000; Harris *et al*. 2018), which follows a rigorous protocol in which skilled volunteers count all birds heard or seen in three distance bands along two 1km transects within a 1km^2^ site on two annual morning visits during the breeding season. The two visits are not designed a priori as replicates, but rather ensure coverage of both early breeding residents and later breeding migrants. The early visit takes place April to mid May, and is followed by a late visit in mid May to June. BBS provides a spatial coverage which is extremely high for a national monitoring scheme (1.10-1.65% of the UK territory for the study period (Harris *et al*. 2018)), and sampling is largely unbiased with respect to habitat types (with the exception of mountainous areas) (Martay *et al*. 2018). The survey follows a stratified random design which aligns coverage with variable volunteer availability. Coverage within strata ranges from 0.1% to 9%. This allows an unbiased assessment of UK-wide and national trends for many common species, but the survey was not designed to allow for inferences at sub-stratum level, or to provide reliable coverage of rare species.

Structured data were supplemented with records from BirdTrack (Newson *et al*. 2016), which is also a citizen science dataset, but with less stringent observation requirements and a wider range of participants than the BBS: last year 2766 volunteers contributed to BBS whereas 6869 individuals submitted at least one BirdTrack list. BirdTrack participants contribute lists of species they have detected during a self-selected time interval spent at a self-selected location. Compared to the BBS there are about three times as many locations (i.e. 1km British National Grid squares) in the UK that have at least one BirdTrack record during the breeding season in recent years (Figure S1). However, the relative density of records for both schemes follows a similar large-scale pattern: Coverage is higher near urban centres and lower in less populated and more mountainous areas (Boersch-Supan *et al*. 2019; Darvill *et al*. 2020). On smaller spatial scales the site-selection biases in BirdTrack are complex. Broadly speaking, sites fall into two clusters: sites that are convenient to access, e.g. in the vicinity of participants’ homes, and more distant sites that offer more diverse bird assemblages (Johnston *et al*. 2020).

We only considered timed complete BirdTrack lists, i.e. lists for which birdwatchers recorded a start and end time and reported that they had listed all detected species. To match the spatial grain and temporal extent of the BBS data we only used lists with a location precision of 1km collected from 01 April to 30 June of each year. The resulting dataset constitutes detection/non-detection data with biases associated with self-selection of sites and visit timings. Finally, we filtered available data to retain only locations which had lists in three or more years, and within a year we randomly sub-sampled lists from locations that had more than 25 visits.

### 2.2 Modelling approach

We used a state-space modelling approach to integrate data from BBS surveys and BirdTrack lists [Isaac *et al*. (2019); Supplementary Materials]. The model assumes an underlying biological process describing species-specific abundances *N*_*j*,*t*_ at a site *j* in every year *t*, and their changes from year to year as a result of individuals that survive and remain at each site, and those that are gained to a site by recruitment or immigration. Following Zipkin *et al*. (2017) we link this process model to the count and detection-nondetection data with a separate observation model for each data source: a dynamic N-mixture model for the count data and a dynamic occupancy model for the detection-nondetection data (Figure S2).

Model parameters were estimated in a Bayesian framework using JAGS via the jagsUI package in R (Plummer 2003; Kellner 2018; R Core Team 2018). Markov-chain Monte Carlo (MCMC) estimation was run on four parallel chains until the Gelman-Rubin convergence diagnostic 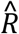 indicated convergence, usually after 10,000-50,000 iterations.

We stratified the modelled sites based on prior information on site occupancy pre-dating the BBS and BirdTrack data from the 1988-1991 Bird Atlas (Gibbons, Reid & Chapman 1993). Sites that fell into occupied tetrads (2km x 2km squares) in the Atlas were assigned a positively-biased normal prior truncated at zero for the initial abundance *N*_+_(5,10), and sites that fell into unoccupied or unsurveyed tetrads were assigned a zero-biased normal prior, truncated at zero, i.e. a half-normal prior *N*_+_(0,10).

Population trajectories from integrated models were compared to relative abundance indices derived from BBS data alone using the standard BBS trend model, a survey weighted count model with fixed additive site and year effects (Freeman *et al*. 2007) (Supplementary Methods) and occupancy indices derived from dynamic occupancy models using BirdTrack data alone which closely mirrored the structure of the integrated model (Kéry *et al*. 2010).

## 3 Case studies

### 3.1 Improving small-area trends

The motivation for this case study was to assess the utility of data integration to draw inferences about population trends areas smaller than a BBS stratum. We chose the Corn Bunting *Miliaria calandra* as the focal species (Figure 2), a lowland farmland bird whose dramatic decline in range and abundance in the UK has made it a red listed species of conservation concern (Eaton *et al*. 2015) and the target of management interventions (Perkins *et al*. 2011). We fitted integrated models for two areas to contrast different levels of recording coverage: the South Downs National Character Area of southern England which has an expanse of c. 1,000km ^2^ (Figure S3), and a similar sized area largely dominated by arable farmland in North East Scotland (Figure S4). The South Downs are close to major conurbations and are well covered by BirdTrack records from recreational birdwatchers; recording in North East Scotland occurs at much lower rates (Figure 2).

**Figure 2:**
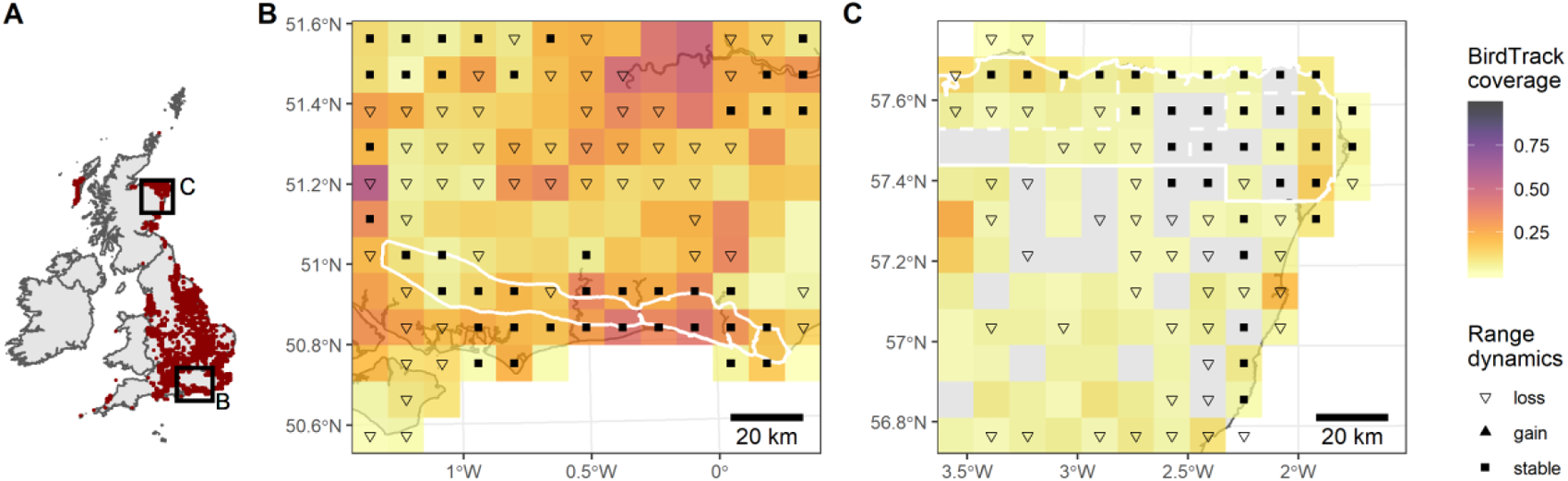
A: The range of Corn Bunting in Great Britain and Ireland 2007-2011. B: The South Downs NCA (white outline) is well covered by BirdTrack observations. C: Farmland in the NE of Scotland is poorly covered by BirdTrack observations. Coloured squares indicate BirdTrack list density. Grey squares lack BirdTrack lists. Symbols indicate Corn Bunting status based on the 2007-2011 Bird Atlas. White outlines show the spatial domain of the integrated models.

#### 3.1.1 Species-specific model details

Survival and colonisation rates were separately estimated within each stratum (occupied, unoccupied, unsurveyed) as a yearly random effect, using an informative Beta prior (mean 0.58, variance 0.24) based on a mark-recapture estimate of survival probability (Luebcke 1977).

#### 3.1.2 Results

Single data source trend models and the integrated trend could be derived for the South Downs NCA. Because of the lower density of records in both schemes in north east Scotland, some or all trend models failed to fit for the two areas with the highest density of BirdTrack lists that were of equal size to the South Downs NCA. Model fitting was successful when using records from an area spanning Elgin to Peterhead, approximately three times the size of the South Downs NCA (Figure 2, Figure S4).

For both areas the trend estimates from the joint model did not differ substantially from occupancy changes derived from BirdTrack or abundance changes derived from BBS alone, respectively. All models for the South Downs NCA showed a range and abundance decline between 2005 and 2011 followed by a period of relative stability (Figure 3A), and the models for North East Scotland yielded highly uncertain abundance and occupancy trends, neither of which provided statistically significant evidence of change since 2005 (Figure 3B).

**Figure 3:**
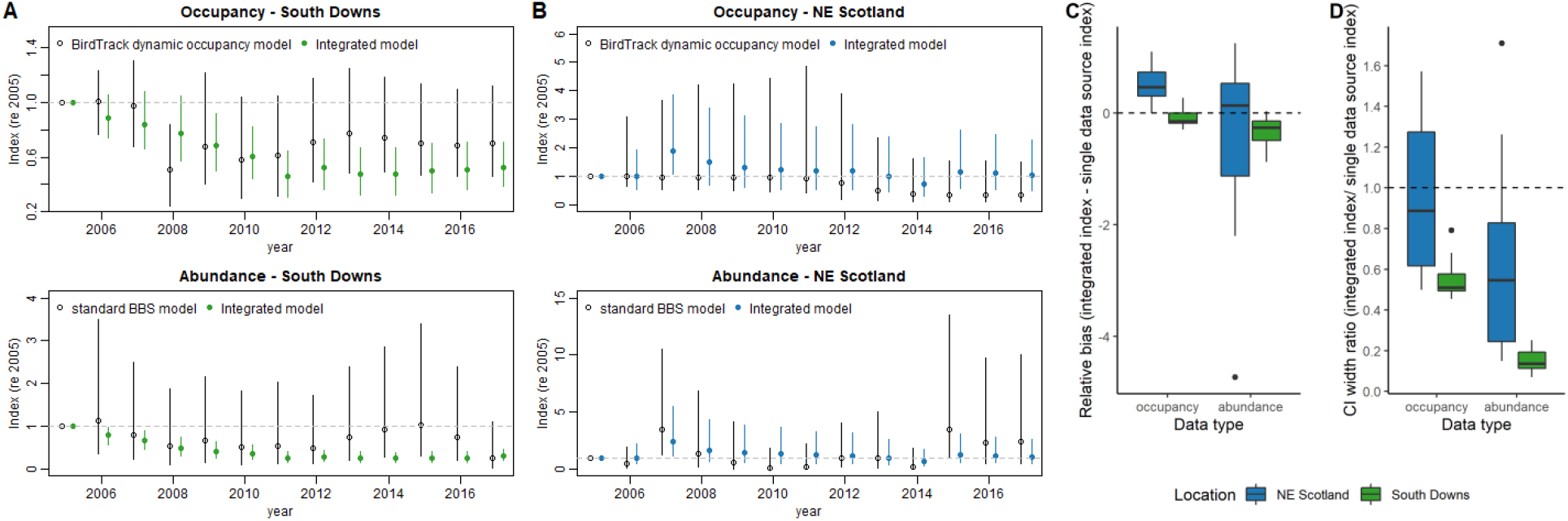
Occupancy and abundance trend estimates for Corn Bunting in the South Downs (A, green) and NE Scotland (B, blue) based on single data sources (BBS *or* BirdTrack; open symbols) and the integrated model (BBS *and* BirdTrack; solid symbols). Error bars show posterior 95% credible intervals. Boxplots aggregate relative bias (C) and precision (D) of annual index values comparing the integrated and single data source models.

The integrated trend was negatively biased compared to both reference trends at both locations, although point estimates for yearly index values from the integrated model were within the credible intervals of each reference trend. In the South Downs credible intervals for the integrated occupancy trend were about 50% as wide as those of the BirdTrack occupancy trend model, and integrated abundance trend credible intervals were even narrower at about 20% of those of the BBS trend (Figure 3C,D). The integrated model predicts a significant decline of Corn Bunting range and abundance in the study area between 2005 and 2011. In contrast, models based on either dataset alone do not allow inferences about population change given the large uncertainty about annual index values. In Scotland inferential gains from data integration were much more modest. The nominal precision of the integrated model results was on par with the BirdTrack occupancy trend, and the credible intervals of the integrated abundance trend were about half as wide as those for the BBS standard model (Figure 3C,D).

Aside from the inferences about range and abundance changes the integrated model also provides estimates of detection parameters such as the influence of time spent recording on the probability of detecting a given species (Figure 4). This is a crucial feature to assess the properties of unstructured observations, as observation protocols for these data are less stringent leading to substantial heterogeneity in observer effort.

**Figure 4:**
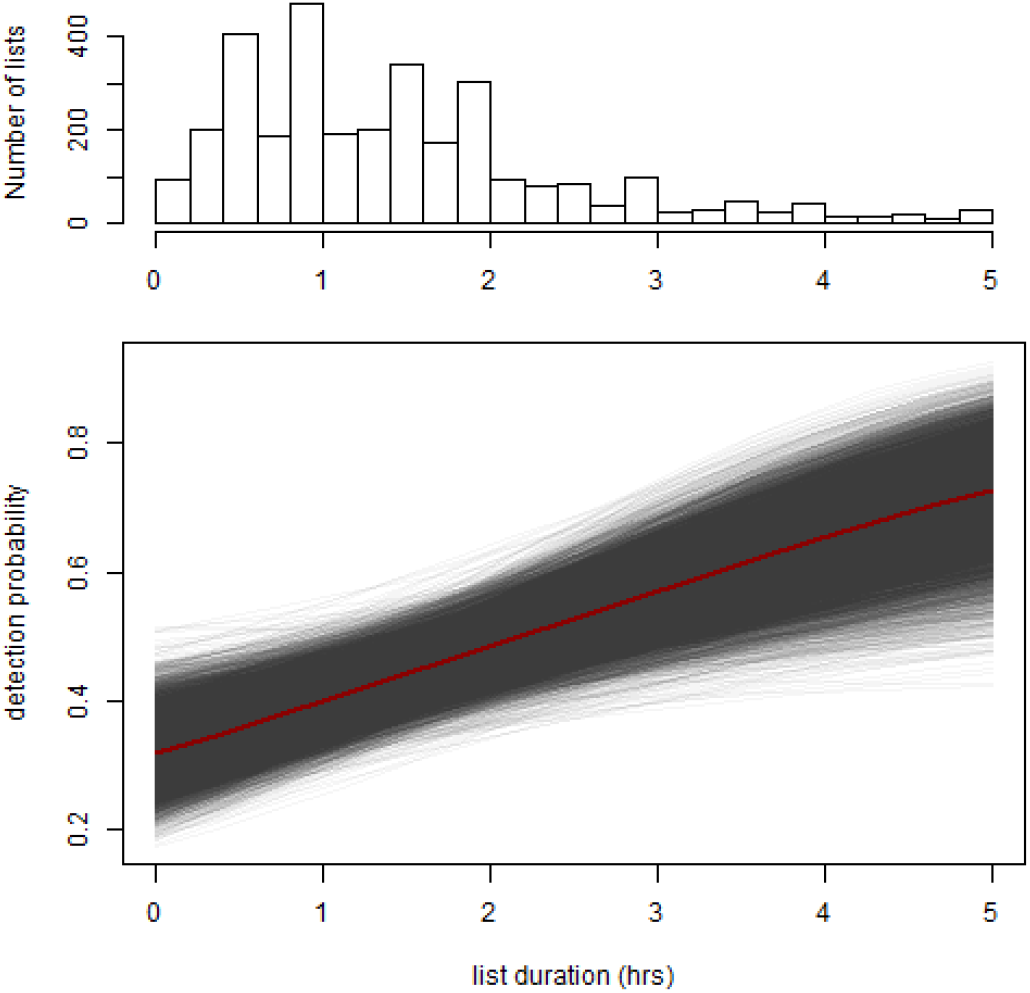
Detection probability of Corn Bunting in the South Downs increases with recording duration. Top: Distribution of BirdTrack list durations (in hours). Bottom: Estimated effect of list duration on the probability of detecting at least one individual. Red line shows median, grey lines show realisations from the detection model posterior.

### 3.2 A rare but widespread species

The second case study assessed the utility of data integration to draw inferences about population trends for a species that is not widespread enough to fulfil the minimum sample size criterion for standard BBS reporting at the country level. We chose the Pied Flycatcher *Ficedula hypoleuca* in Wales as the focal species (Figure S5). It is a migratory woodland bird with a distribution restricted to upland deciduous woods in parts of western and northern Britain. It is red listed, both in Wales and UK wide, due to its breeding population decline over the last 25 years (Eaton *et al*. 2015; Johnstone & Bladwell 2016).

#### 3.2.1 Results

In the relative comparison the occupancy change estimates from the joint model did not differ substantially from occupancy changes derived from BirdTrack alone (Figure 5).

**Figure 5:**
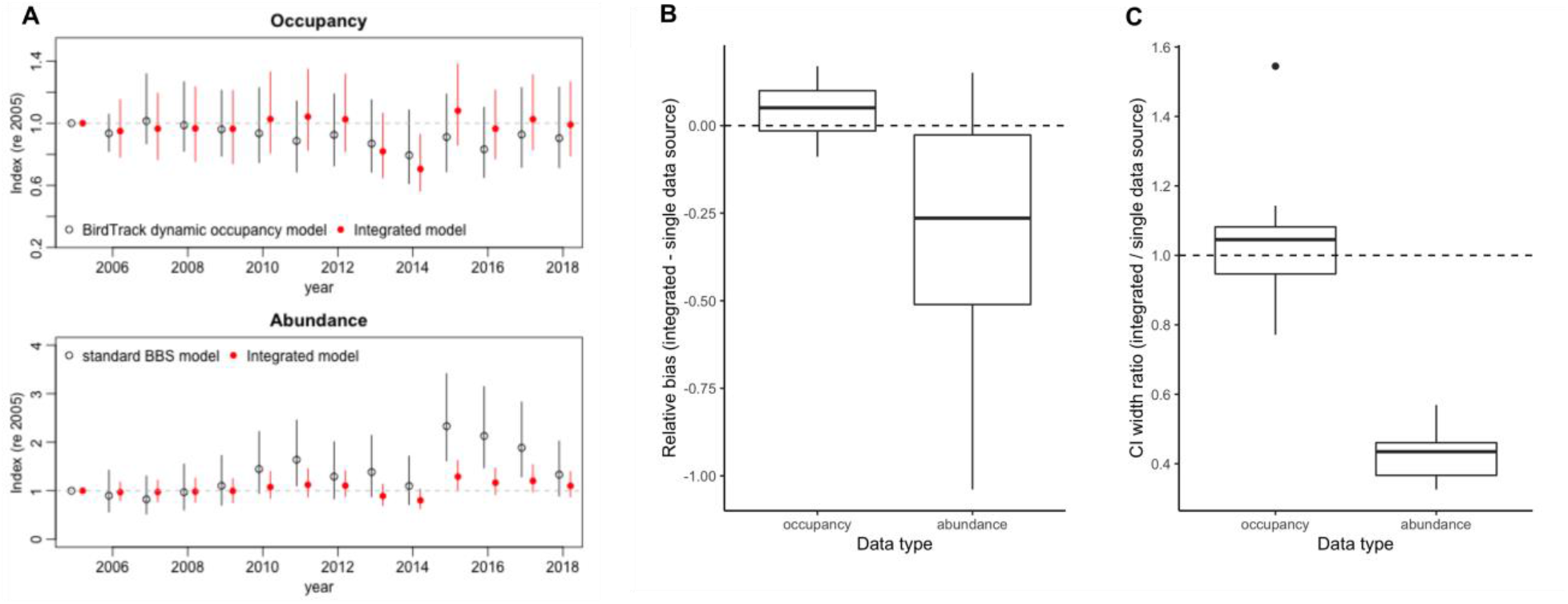
A: Occupancy and abundance trend estimates for Pied Flycatcher in Wales based on single data sources (BBS *or* BirdTrack; open symbols) and the integrated model (BBS *and* BirdTrack; red solid symbols). Error bars show posterior 95% credible intervals. B,C: Relative bias and precision of annual index values comparing the integrated and single data source models..

The integrated abundance trend was more precise with credible intervals about 40% the width of those of the BBS trend, however, the integrated trend was negatively biased compared to the BBS trend (Figure 5 B,C). This bias was strong enough in recent years to result in diverging inferences between the two models. The BBS model indicates that the Welsh Pied Flycatcher population is growing, with significant gains in 2011 and 2015-2017 compared to the reference year 2005. Population fluctuations indicated by the integrated trend model followed a similar pattern of gains and losses but with overall smaller magnitudes of change, resulting in a prediction of a stable population with no statistically significant gains or losses in any year since 2005 (Figure 5 A). Despite their discrepancy both of these findings are more optimistic than trends observed for other humid-zone migrants in England (Morrison *et al*. 2013).

## 4 Discussion

Integrated models of BBS and BirdTrack data provided realistic estimates of the regional population trajectories of both bird species. In all cases the integrated trends had higher nominal precision for the abundance trajectory compared to models based on structured count data alone. Integrated occupancy trends were at least as precise as, or more precise than occupancy trends based on unstructured detection-nondetection data alone. Bias in the trends for both species was harder to assess. Population trajectories followed similar shapes, but overall effect sizes differed between models. Further work is required to validate integrated trend models against existing independent survey data, and to develop cross-validation strategies for evaluating integrated models in the absence of independent reference data. The availability of independent reference data is limited to a small number of species and regions (e.g. Stevens, Murn & Hennessey (2019), Stevens, Murn & Hennessey (2020)). In part this is due to the relative recent development of BirdTrack, which means that, as yet, temporal overlap with several established national surveys for rare bird species (e.g. Cirl Bunting (Stanbury *et al*. 2010), Dotterel (Hayhow *et al*. 2015), Hen Harrier (Wotton *et al*. 2018)) is insufficient for formal comparisons - a situation that should improve in the future.

Although the BBS collects detectability data using distance sampling, this information is currently not included in the calculation of routine BBS trends, as the effects of heterogeneous detection on trend estimates are deemed small (Newson *et al*. 2013). However, this assumption is less likely to hold for rare species. The integrated model therefore attempted to capture observation uncertainty in structured data using an N-mixture model. Although this type of observation model has been shown to be robust under certain field conditions (Bötsch, Jenni & Kéry 2019), N-mixture models are known to be sensitive to violations of their assumptions, including the closure assumption (i.e. that there is no change in occupancy between survey visits) (Barker *et al*. 2018). Given the two BBS visits are not designed as replicates this assumption only holds for species with relatively unchanged detectability within the survey window. Using the distance sampling data to model detectability in the BBS data would be preferable (Farr, Green & Zipkin 2020), however, the corresponding observation model is computationally much more demanding than the N-mixture model. Computational effort for the estimation of integrated model parameters was high with the larger case study requiring model fitting times in the order of 5-24 hours on dedicated scientific computing hardware with Intel Xeon E5 processors and ample memory. This makes model development and checking slow and may limit the roll-out of this model type for routine reporting across many species and regions.

The expected gains from data integration will vary both depending on the target species and target area. Target areas are an important consideration because sampling coverage for both structured and unstructured data are not evenly distributed (Figure 6). The modelling approach used for the Corn Bunting case study performed well in the South Downs because this area has good BBS coverage and exceptional BirdTrack coverage. In North East Scotland low coverage from BBS and very low coverage from BirdTrack made it impossible to fit some or all trend models for an area equivalent to the South Downs NCA (c. 1,000 km ^2^). Model fitting did succeed when increasing the spatial domain to about 3,000km ^2^, but even then gains in precision from the integrated approach were modest. In fact, for much of Scotland the target species range does not overlap with the distribution of opportunistic sampling effort (Figure 2), making it impossible to gain information from opportunistic observations e.g. about the efficacy of agri-environment schemes and indicating the lower limit at which such data might inform policy.

Similarly, the modelling strategy used here would likely not provide gains for country-level trends in Northern Ireland at the current level of BirdTrack coverage (Figure S1). Integration of opportunistic records is thus not a silver bullet for closing gaps in biodiversity monitoring on sub-national scales. At current levels of recording in the UK precision gains in bird trends from data integration can be expected at most NUTS 1 units (e.g. countries, statistical regions of England), densely settled NUTS 2 units (e.g. counties) or similarly sized landscape units such as NCAs, but likely only few NUTS 3 units (e.g. unitary authorities, districts, council areas).

In summary, we demonstrate that integration of structured and unstructured biodiversity records is in principle feasible for trend reporting at national and sub-national scale. Given the growing popularity of recreational biodiversity recording, opportunistic records are available in many countries which also maintain structured survey schemes, making our approach transferable beyond the UK and to non-avian taxa. However our findings also highlight that addressing monitoring gaps at these scales can not be solved with statistical models alone, but requires a careful consideration of the most promising survey approaches: In densely populated areas existing opportunistic citizen science schemes may provide a relatively easy solution to fill information gaps, but elsewhere information gains require steering the observation efforts in both opportunistic (see e.g. Callaghan *et al*. (2019b), Callaghan *et al*. (2019a)) and structured schemes, as is done e.g. for the BBS by targeted efforts to increase observer coverage of mountainous survey strata (Darvill *et al*. 2020). Birds are disproportionally well covered by both structured and unstructured schemes within the UK and globally (Amano *et al*. 2016; Sorte & Somveille 2020). Given the generally lower coverage of non-avian taxa by structured surveys, the potential for relative information gain from opportunistic schemes is expected to be much larger. At the same time, our findings imply that scheme design considerations are likely even more important for these taxa to ensure that spatially biased and/or heterogenous coverage from opportunistic observations at national and sub-national scales does not affect the representativity of derived trends.

## 5 Author contributions

PHBS and RAR conceived the study; PHBS led data analysis and writing with input from RAR. Both contributed critically to the manuscript and gave final approval for publication.

## 6 Data availability statement

The BTO is committed to making data and information readily available. Data underlying this study are available by making a direct request to, or through our website https://www.bto.org/our-science/data/data-request-system.

## 7 Acknowledgements

We thank the thousands of volunteers who contribute records to BirdTrack and the BBS and past and present survey organizers and staff, particularly Sarah Harris, Scott Mayson, Steve Pritchard, and Andy Musgrove. BirdTrack is operated by the BTO, and supported by the RSPB, BirdWatch Ireland, Scottish Ornithologists’ Club, and the Welsh Ornithological Society. The BTO/JNCC/RSPB Breeding Bird Survey is a partnership jointly funded by the BTO, RSPB, and JNCC. This study was part funded by JNCC through the Terrestrial Surveillance Development and Analysis partnership. Computations for this study used JASMIN, the UK’s collaborative data analysis environment (http://jasmin.ac.uk). We thank David Allen, Stephen Baillie, Alison Eyres, Simon Gillings, and Paula Lightfoot for comments on earlier versions of this manuscript.

## 9 Supplementary Materials

**Figure S1:**
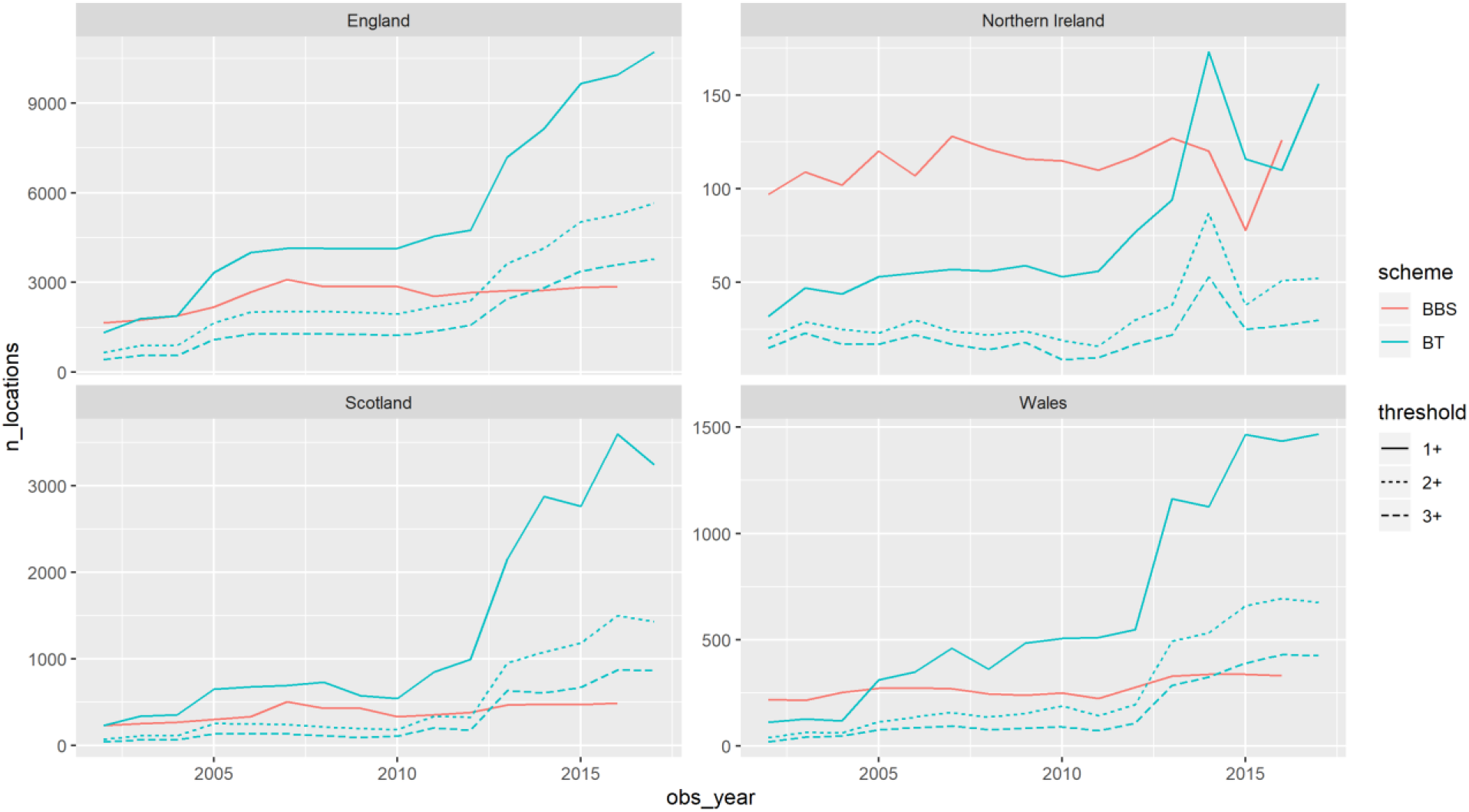
Opportunistic sampling coverage (measured here as annual locations with lists, solid blue line) has greatly increased since the inception of the BirdTrack scheme and now exceeds the number of BBS plots (solid red line) in all four countries. Revisits of locations by BirdTrack participants are relatively rare, however, with only about half of the sites having lists in two or more years (dotted blue line), and even fewer in three or more years (dashed blue line).

### 9.1 Integrated Model details

We used a state-space modelling approach which models the latent biological processes of population persistence and growth and links them to the observed data using an observation model that accounts for imperfect detection to integrate data from BBS surveys and BirdTrack lists (Figure S2:). This integrated model is compared to trends derived from each of the two data sources separately. For the structured BBS data we use the statistical model that is used in the official reporting of this survey (Freeman *et al*. 2007) as a reference model.

This is a simple trend model in that it relies on the randomised nature of the survey and does not explicitly model the observation process. For the unstructured BirdTrack data the reference model we use is a dynamic occupancy model which explicitly models the observation process (van Strien *et al*. 2013). This class of models has been shown to be reasonably robust for opportunistic biological records (Kéry *et al*. 2010; van Strien *et al*. 2013), unlike simpler trend models (Boersch-Supan *et al*. 2019).

**Figure S2:**
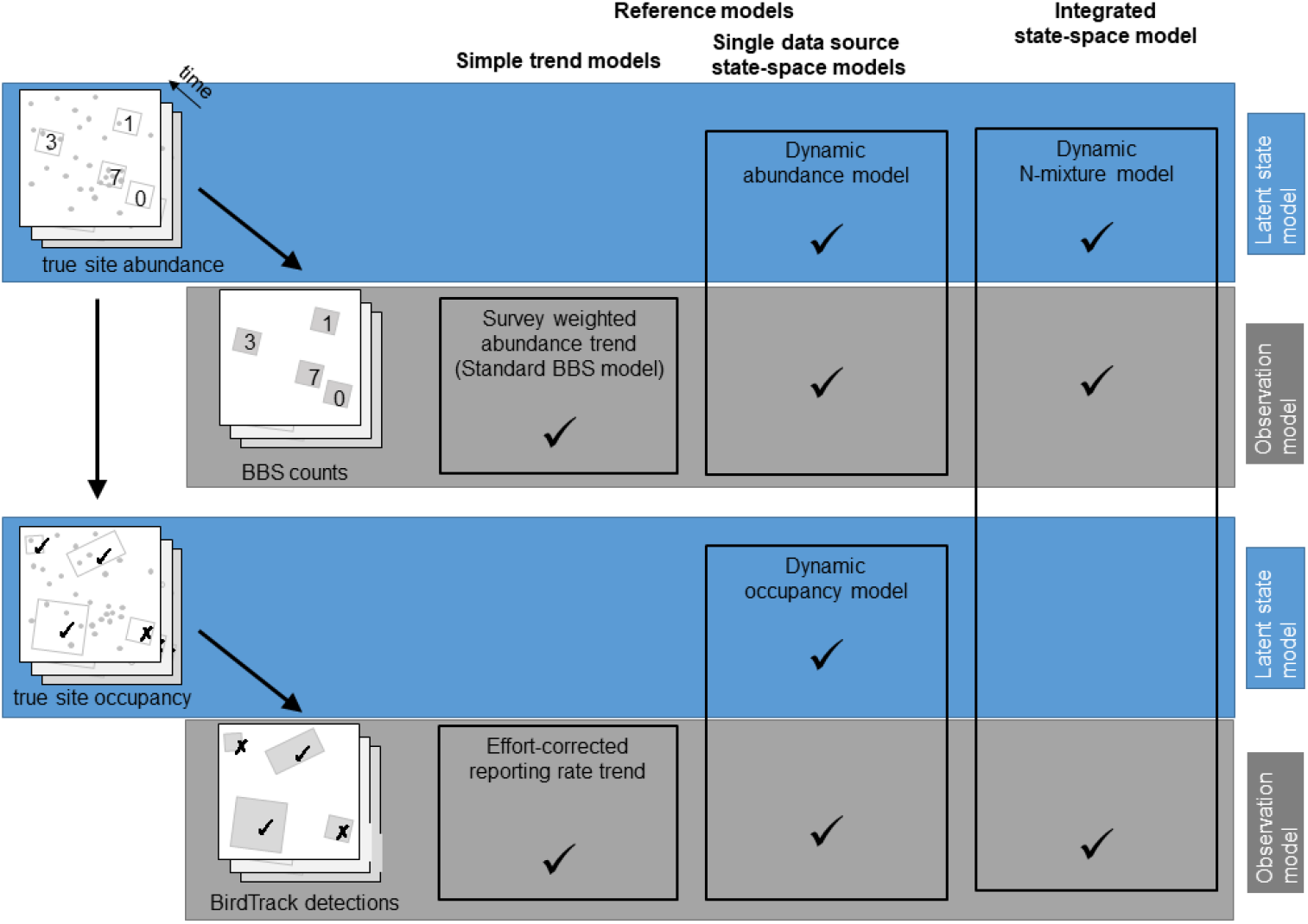
An overview of the modelling approaches used in his study. Reference models cover a single dataset at a time and serve as a comparison with the integrated model. There are two types of reference models, simple regression models of the observations and hierarchical models that explicitly separate the biological state process (i.e. the latent population dynamics) from the observation process. Data integration relies on this conceptual separation and link the two different data sources via a common process model.

#### 9.1.1 Biological process model

The state process for this model assumes there is a population abundance *N*_*j*,*t*_ at a site *j* in every time step *t*, which is imperfectly observed. *N*_*j*,*t*_ changes between successive timesteps as a result of individuals that survive and remain at each site *S*_*j*,*t*_, and those that are gained to a site by recruitment or immigration *G*_*j*,*t*_. These sub-processes are expressed as

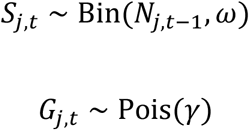

where *ω* is the apparent annual survival probability of individuals, and *γ* is the expected number of individuals that are gained at site *j* by recruitment or immigration between *t* − 1 and *t*.

For every time step *t* > 1 the total population abundance at site *j* is

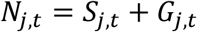

For the first year (*t* = 1), the state process is initialized by modelling abundance at each site according to a Poisson distribution with an expected count *λ*

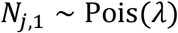

From the state model we can further derive the colonisation probability *ϕ*_*j*,*t*_ of an unoccupied site as

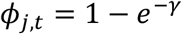

as well as the extinction probability *ϵ*_*j*,*t*_ as

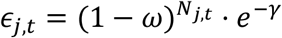

#### 9.1.2 Observation model

The true population abundance *N*_*j*,*t*_ in the survey area was linked to the data according to two sampling processes, counts of individuals in the case of BBS data and detection of at least one individual or non-detection in the case of BirdTrack lists. In both cases repeat visits to a site between April and the end of June were treated as replicates, implying the assumption of a closed population over this period, which we deemed reasonable for the species covered.

We assumed detection was imperfect for both sampling approaches, i.e. for count data the number of individuals encountered during a survey visit *n*_*j*,*t*,*k*_ ≤ *N*_*j*,*t*_ and similarly, an occurrence record *y*_*j*,*t*,*k*_ could be a nondetection if none of the *N*_*j*,*t*_ individuals was seen or heard during a site visit. We modelled the count data as arising from a binomial process

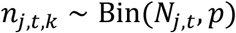

with an individual detection probability *p*. Detection-nondetection data were modelled as arising from a Bernoulli trial

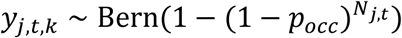

with a separate detection probability *p*_*occ*_, to take into account potential differences in survey methodology and/or observer skill between BBS and BirdTrack records.

### 9.2 Reference Trends

#### 9.2.1 BBS abundance trends

Abundance models for BBS data followed the Poisson GLM approach employed in the official BBS trend production (Freeman *et al*. 2007), which models the mean local count *λ*_*it*_ at site *i* and year *t* based on the observed maximum counts *y*_*obs*,*it*_ across the two survey visits as a function of fixed additive site and year effects *β*_*i*_ and *β*_*t*_, respectively.

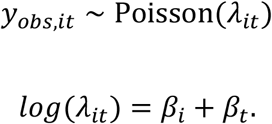

Abundance indices are derived from the conditional year effects *β*_*t*_. For BBS data we further used sampling weights – equal to the inverse inclusion probability of a site within a stratum for a given year – to account for uneven monitoring coverage among BBS survey strata.

Parameter inference was conducted in a Bayesian framework using a weighted likelihood approach as implemented in the brms package (Bürkner 2018), rather than following the bootstrapping approach of Freeman *et al*. (2007).

#### 9.2.2 BirdTrack Dynamic Occupancy models

Occupancy trends from BirdTrack data were modelled using a dynamic site occupancy model (Kéry *et al*. 2010; van Strien *et al*. 2013) which closely mirrored the structure of the integrated model. The latent state *N*_*j*,*t*_ becomes a binary indicator of occupancy in these models, and initial occupancy *N*_*j*.1_ and the immigration/recruitment process *G*_*j*,*t*_ were modelled as a Bernoulli processes.

### 9.3 Data details

**Figure S3:**
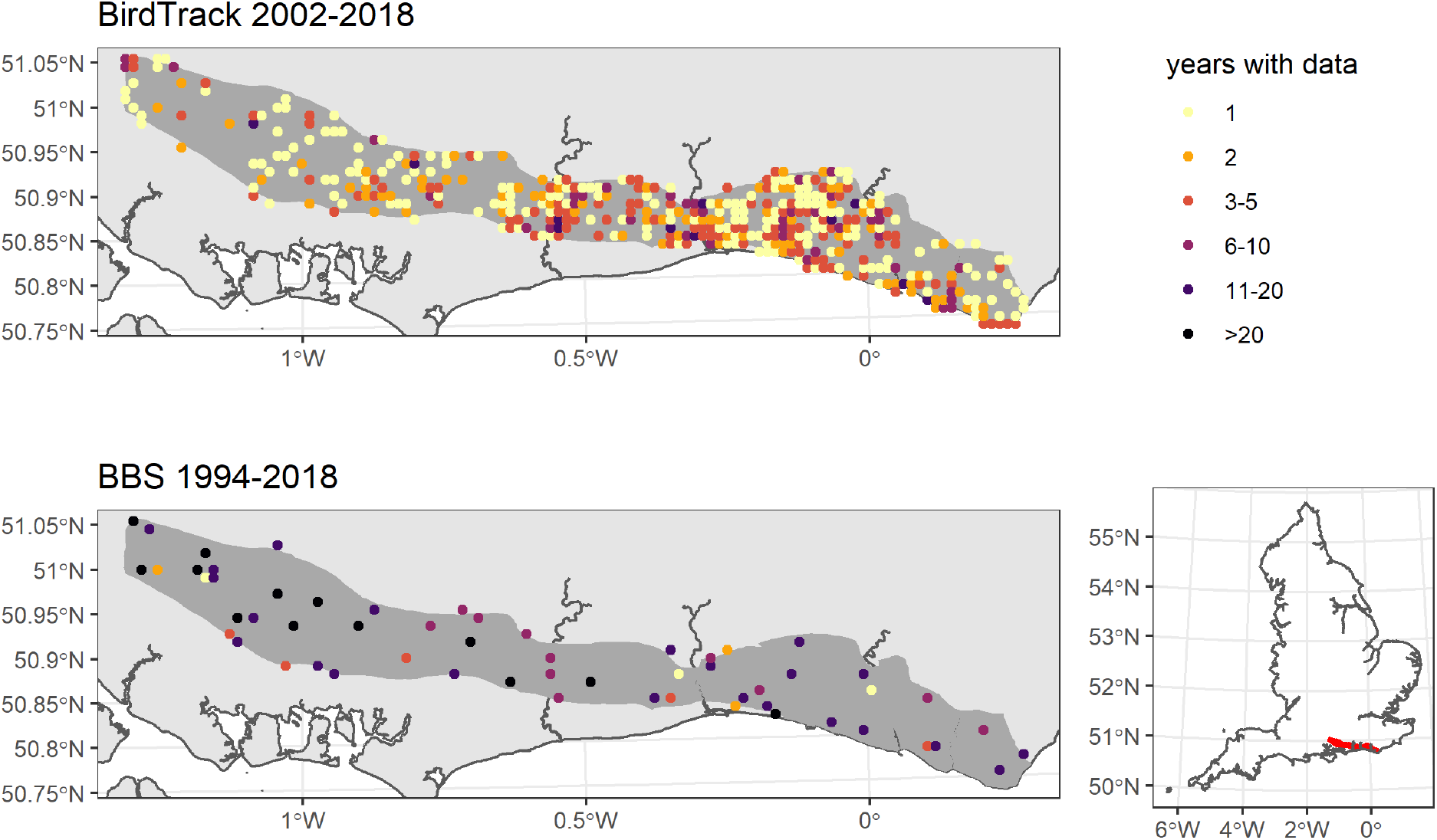
The study area for the South Downs Corn Bunting case study and locations of BBS surveys and complete BirdTrack lists.

**Figure S4:**
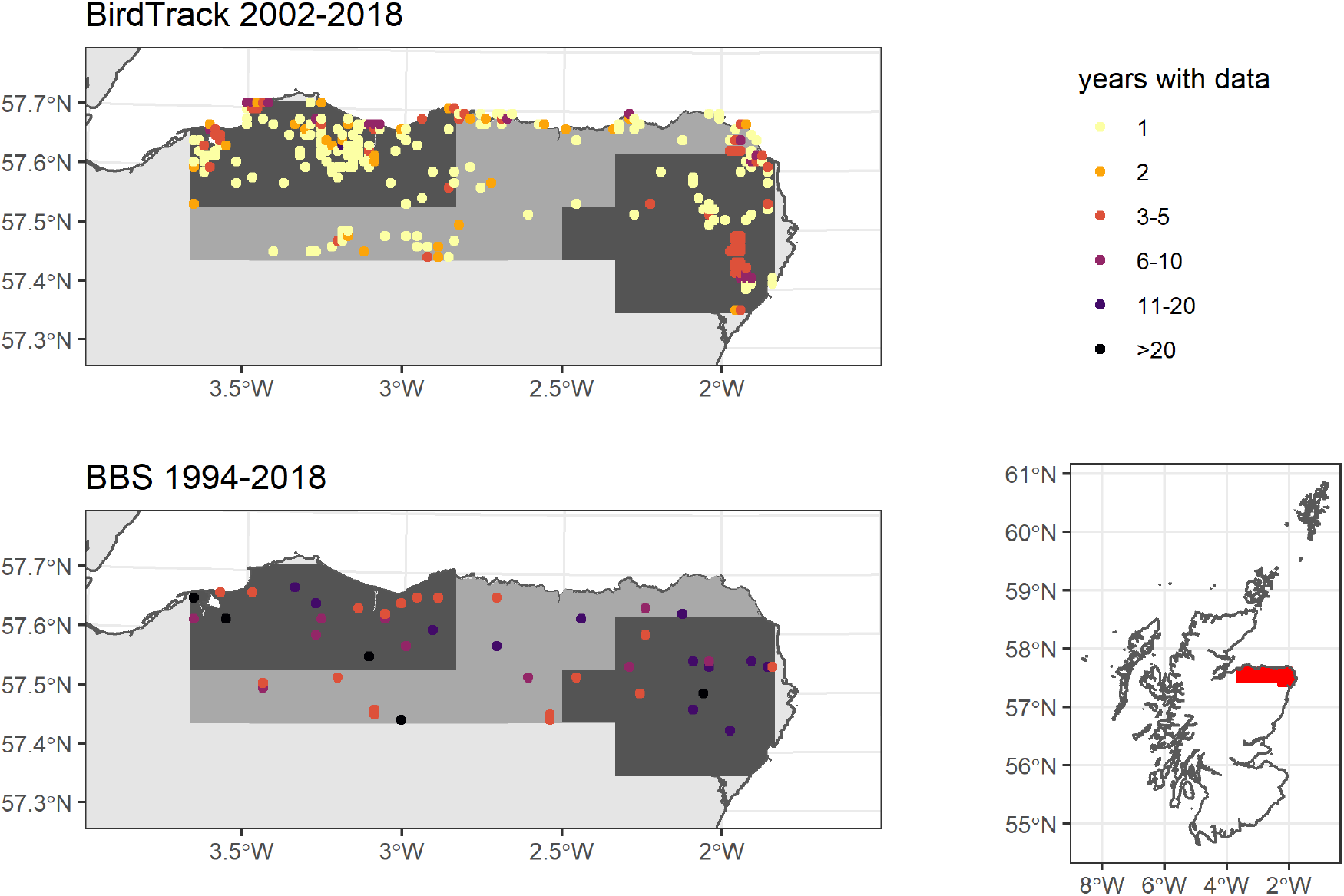
The study area for the Scottish Corn Bunting case study and locations of BBS surveys and complete BirdTrack lists. Both dark shaded areas are approximately the same size as the South Downs NCA (Fig. S3), but data were to sparse in both to fit the integrated and/or references models. Model fitting succeeded when using records from the dark and medium grey shaded areas.

**Figure S5:**
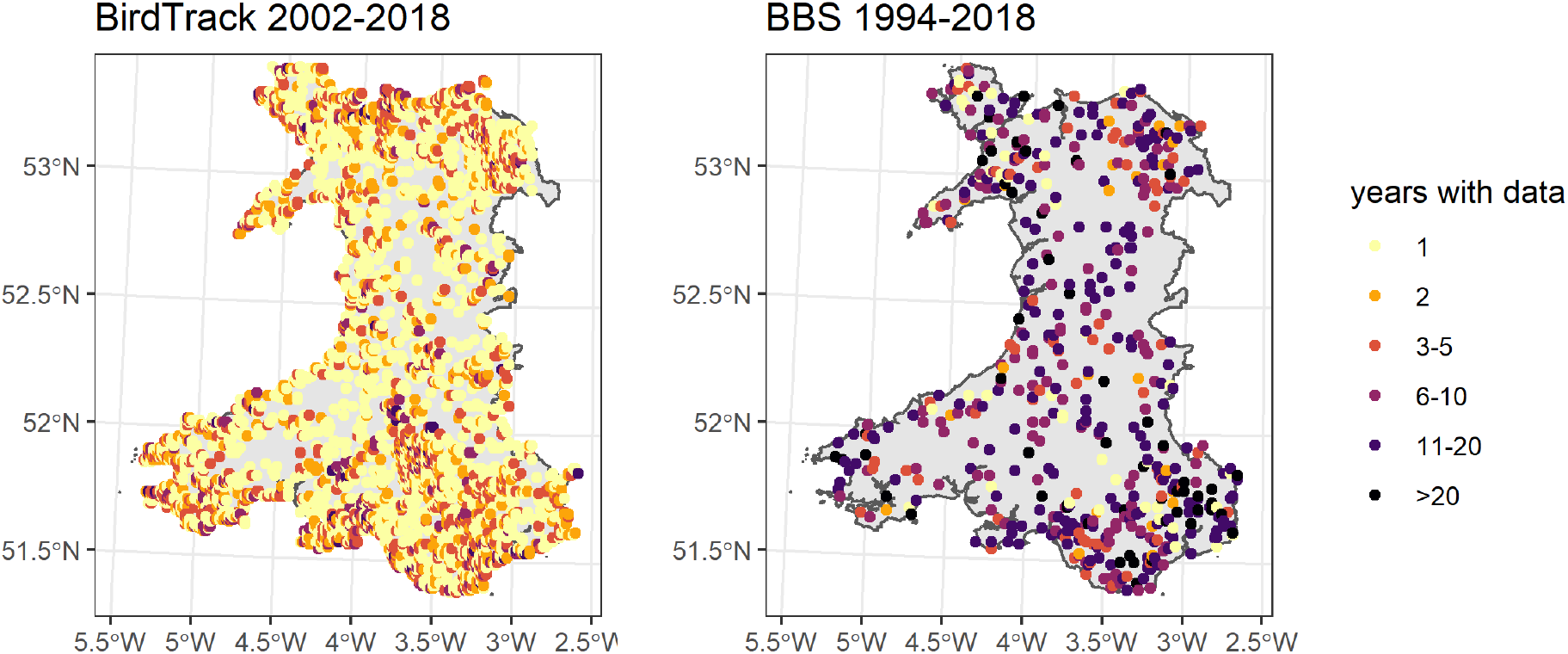
Top: The study area for the Pied Flycatcher case study and locations of BBS surveys (n=519) and locations of complete BirdTrack lists (n=1034). Bottom: Time-series of the number of locations with BBS surveys and complete BirdTrack lists (solid) and locations with positive detections of the target species (dashed).

### 9.4 Model code examples

#### 9.4.1 BBS survey weighted trend models

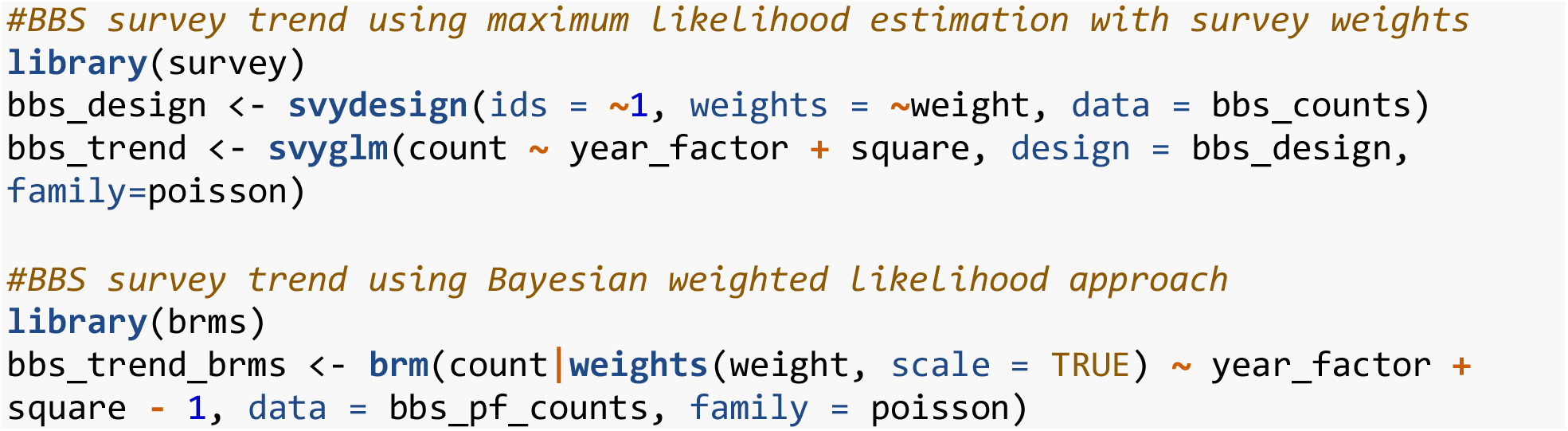

#### 9.4.2 BirdTrack dynamic occupancy model JAGS code

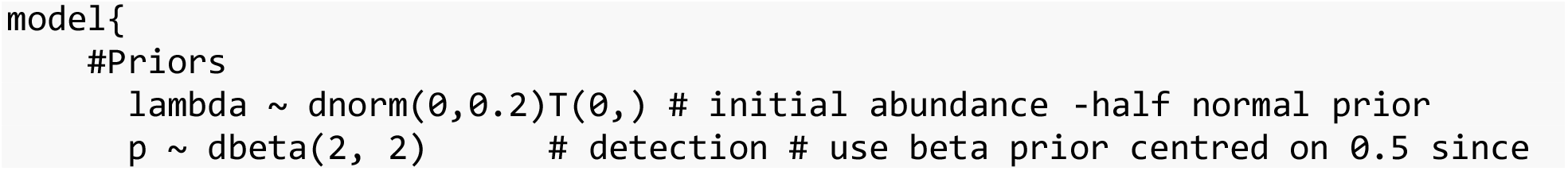
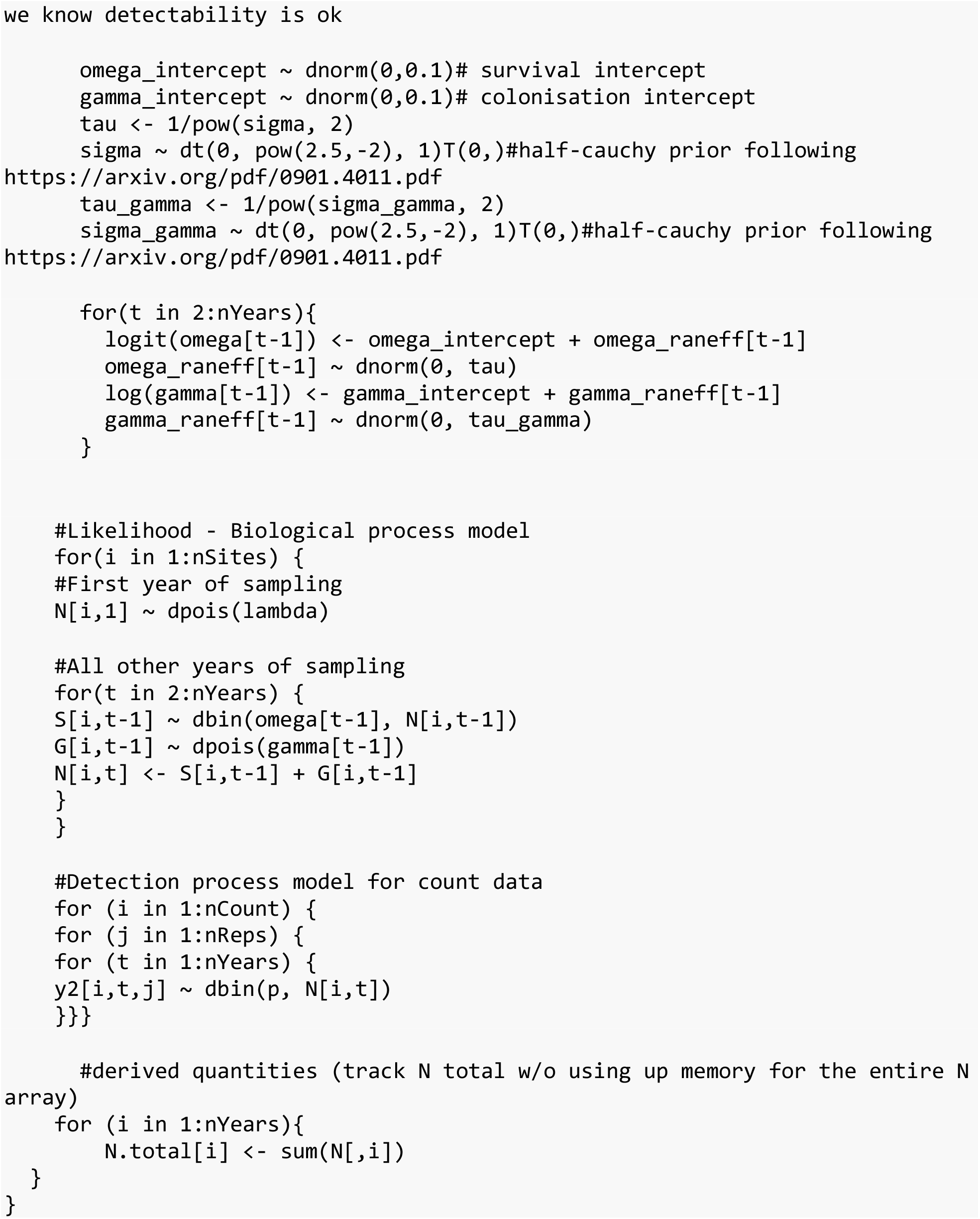

#### 9.4.3 Joint model JAGS code for corn bunting case study

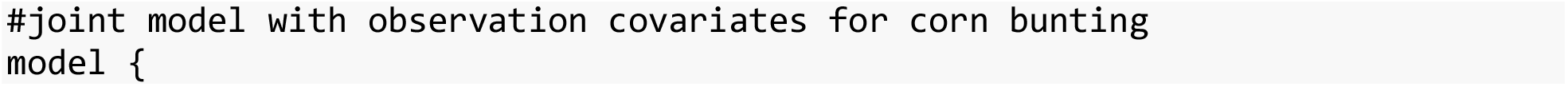
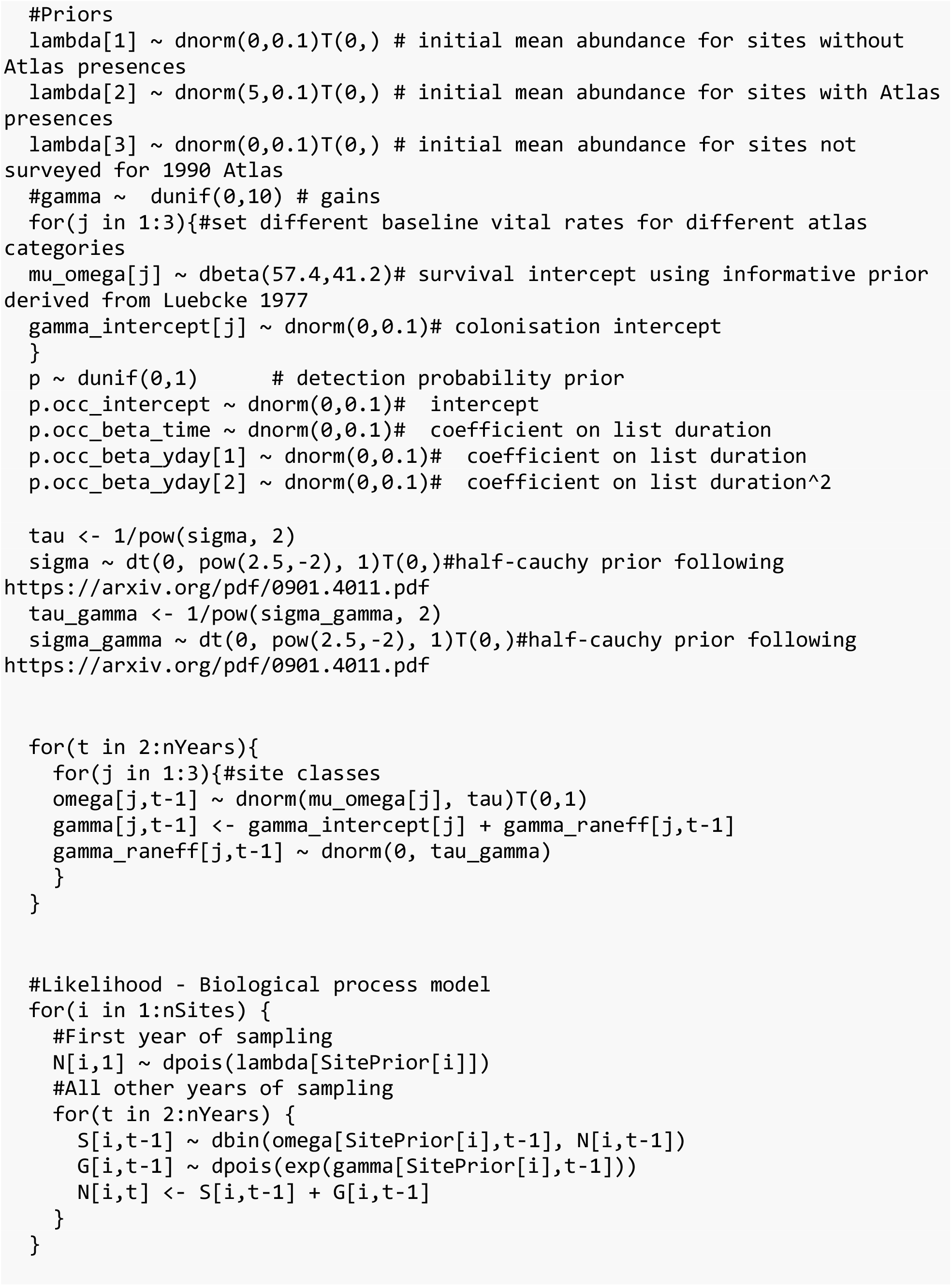
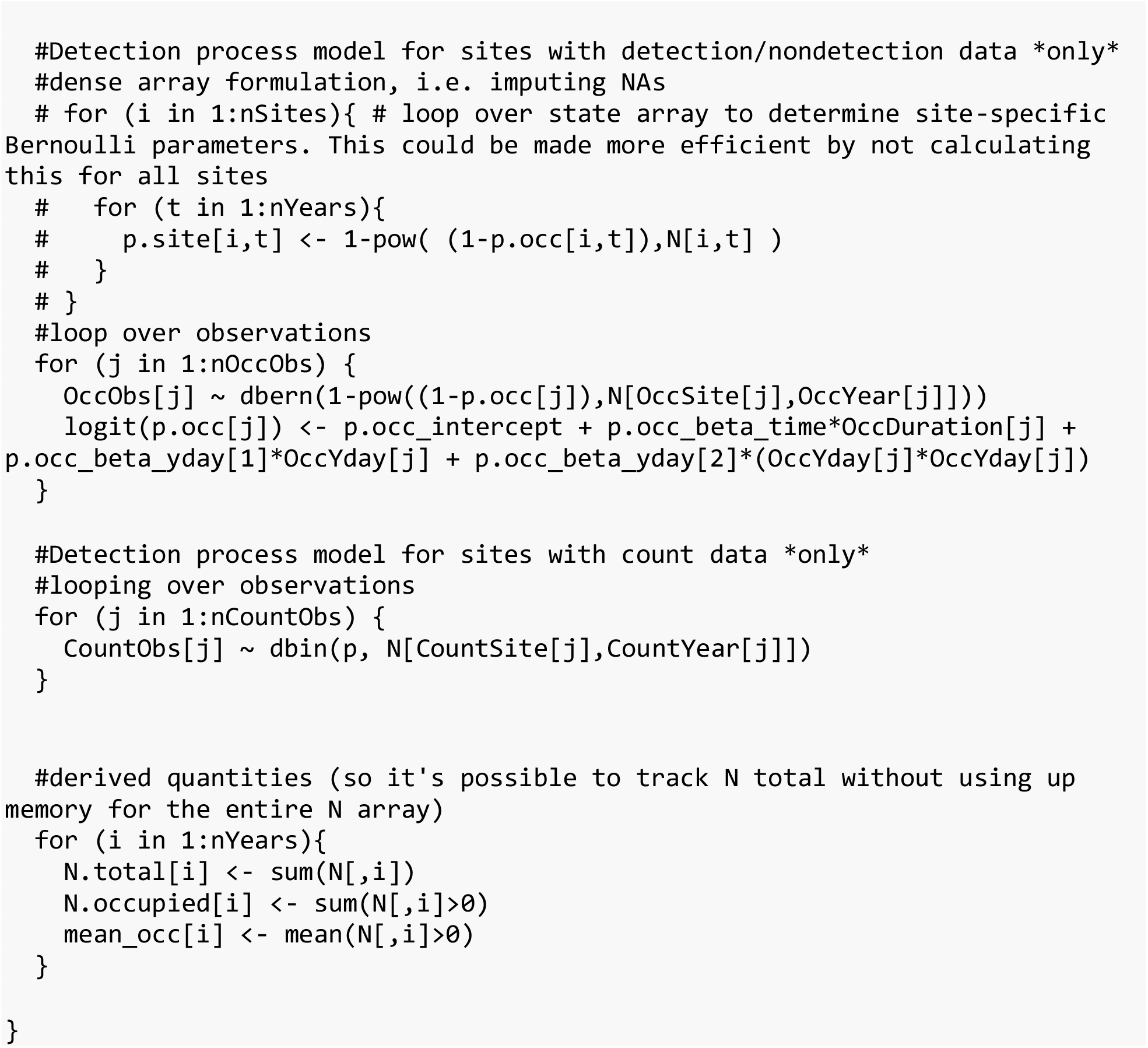

## References

Aceves-Bueno, E., Adeleye, A.S., Feraud, M., Huang, Y., Tao, M., Yang, Y. & Anderson, S.E. (2017) The accuracy of citizen science data: A quantitative review. The Bulletin of the Ecological Society of America, 98, 278–290.

Amano, T., Lamming, J.D. & Sutherland, W.J. (2016) Spatial gaps in global biodiversity information and the role of citizen science. Bioscience, 66, 393–400.

Baillie, S.R., Balmer, D.E., Downie, I.S. & Wright, K.H. (2006) Migration watch: An internet survey to monitor spring migration in Britain and Ireland. Journal of Ornithology, 147, 254–259.

Bainbridge, I. (2014) How can ecologists make conservation policy more evidence based? Ideas and examples from a devolved perspective. Journal of Applied Ecology, 51, 1153–1158.

Barker, R.J., Schofield, M.R., Link, W.A. & Sauer, J.R. (2018) On the reliability of n-mixture models for count data. Biometrics, 74, 369–377.

Bayraktarov, E., Ehmke, G., O’Connor, J., Burns, E.L., Nguyen, H.A., McRae, L., Possingham, H.P. & Lindenmayer, D.B. (2018) Do big unstructured biodiversity data mean more knowledge? Frontiers in Ecology and Evolution, 6, 239.

Birdlife International. (2004) Birds in Europe: Population Estimates, Trends and Conservation Status. Birdlife International, Cambridge, UK.

Boersch-Supan, P.H., Trask, A.E. & Baillie, S.R. (2019) Robustness of simple avian population trend models for semi-structured citizen science data is species-dependent. Biological Conservation, 240, 108286.

Bötsch, Y., Jenni, L. & Kéry, M. (2019) Field evaluation of abundance estimates under binomial and multinomial n-mixture models. Ibis.

Burns, F., Eaton, M., Hayhow, D., Outhwaite, C., Al Fulaij, N., August, T., Boughey, K., Brereton, T., Brown, A., Bullock, D. & others. (2018) An assessment of the state of nature in the united kingdom: A review of findings, methods and impact. Ecological indicators, 94, 226–236.

Bürkner, P.-C. (2018) Advanced Bayesian multilevel modeling with the R package brms. The R Journal, 10, 395–411.

Callaghan, C.T., Poore, A.G., Major, R.E., Rowley, J.J. & Cornwell, W.K. (2019a) Optimizing future biodiversity sampling by citizen scientists. Proceedings of the Royal Society B, 286, 20191487.

Callaghan, C.T., Rowley, J.J., Cornwell, W.K., Poore, A.G. & Major, R.E. (2019b) Improving big citizen science data: Moving beyond haphazard sampling. PLoS biology, 17, e3000357.

Darvill, B., Harris, S., Martay, B., Wilson, M. & Gillings, S. (2020) Delivering robust population trends for scotland’s widespread breeding birds. Scottish Birds, 40, 250–257.

Dickinson, J.L., Zuckerberg, B. & Bonter, D.N. (2010) Citizen science as an ecological research tool: Challenges and benefits. Annual Review of Ecology, Evolution, and Systematics, 41, 149– 172.

Eaton, M., Aebischer, N., Brown, A., Hearn, R., Lock, L., Musgrove, A., Noble, D., Stroud, D. & Gregory, R. (2015) Birds of conservation concern 4:The population status of birds in the uk, channel islands and isle of man. British Birds, 108, 708–746.

Farr, M., Green, D. & Zipkin, E. (2020) Integrating distance sampling and presence-only data to estimate species abundance. Ecology, in press.

Fithian, W., Elith, J., Hastie, T. & Keith, D.A. (2015) Bias correction in species distribution models: Pooling survey and collection data for multiple species. Methods in Ecology and Evolution, 6, 424–438.

Freeman, S.N., Noble, D.G., Newson, S.E. & Baillie, S.R. (2007) Modelling population changes using data from different surveys: The common birds census and the breeding bird survey. Bird Study, 54, 61–72.

Gardiner, M.M., Allee, L.L., Brown, P.M., Losey, J.E., Roy, H.E. & Smyth, R.R. (2012) Lessons from lady beetles: Accuracy of monitoring data from us and uk citizen-science programs. Frontiers in Ecology and the Environment, 10, 471–476.

Gibbons, D.W., Reid, J.B. & Chapman, R.A. (1993) The New Atlas of Breeding Birds in Britain and Ireland: 1988-1991. T & AD Poyser, London.

Greenwood, J.J., Baillie, S.R., Gregory, R.D., Peach, W.J. & Fuller, R.J. (1995) Some new approaches to conservation monitoring of british breeding birds. Ibis, 137, S16–S28.

Gregory, R., Baillie, S. & Bashford, R. (2000) Monitoring breeding birds in the United Kingdom. Bird Census News, 13, 101–112.

Harris, S., Massimino, D., Gillings, S., Eaton, M., Noble, D., Balmer, D., Procter, D., Pearce-Higgins, J. & Woodcock, P. (2018) The Breeding Bird Survey 2017. British Trust for Ornithology, Thetford.

Hayhow, D., Eaton, M., Stanbury, A., Burns, F., Kirby, W., Bailey, N., Beckmann, B., Bedford, J., Boersch-Supan, P., Coomber, F. & others. (2019) State of Nature 2019. State of Nature Partnership.

Hayhow, D.B., Ewing, S.R., Baxter, A., Douse, A., Stanbury, A., Whitfield, D.P. & Eaton, M.A. (2015) Changes in the abundance and distribution of a montane specialist bird, the dotterel charadrius morinellus, in the uk over 25 years. Bird Study, 62, 443–456.

Horns, J.J., Adler, F.R. & Şekercioğlu, Ç.H. (2018) Using opportunistic citizen science data to estimate avian population trends. Biological Conservation, 221, 151–159.

Isaac, N.J. & Pocock, M.J. (2015) Bias and information in biological records. Biological Journal of the Linnean Society, 115, 522–531.

Isaac, N.J., Jarzyna, M.A., Keil, P., Dambly, L.I., Boersch-Supan, P.H., Browning, E., Freeman, S.N., Golding, N., Guillera-Arroita, G., Henrys, P.A., Jarvis, S., Lahoz-Monfort, J., Pagel, J., Pescott, O.L., Schmucki, R., Simmonds, E. & O’Hara, R.B. (2019) Data integration for large scale models of species distributions. Trends in Ecology and Evolution.

Isaac, N.J., van Strien, A.J., August, T.A., Zeeuw, M.P. de & Roy, D.B. (2014) Statistics for citizen science: Extracting signals of change from noisy ecological data. Methods in Ecology and Evolution, 5, 1052–1060.

Johnston, A., Fink, D., Hochachka, W.M. & Kelling, S. (2018) Estimates of observer expertise improve species distributions from citizen science data. Methods in Ecology and Evolution, 9, 88–97.

Johnston, A., Fink, D., Reynolds, M.D., Hochachka, W.M., Sullivan, B.L., Bruns, N.E., Hallstein, E., Merrifield, M.S., Matsumoto, S. & Kelling, S. (2015) Abundance models improve spatial and temporal prioritization of conservation resources. Ecological Applications, 25, 1749– 1756.

Johnston, A., Hochachka, W., Strimas-Mackey, M., Gutierrez, V.R., Robinson, O., Miller, E., Auer, T., Kelling, S. & Fink, D. (2019) Best practices for making reliable inferences from citizen science data: Case study using eBird to estimate species distributions. bioRxiv, 574392.

Johnston, A., Moran, N., Musgrove, A., Fink, D. & Baillie, S. (2020) Estimating species distributions from spatially biased citizen science data. Ecological Modelling.

Johnstone, I. & Bladwell, S. (2016) Birds of conservation concern in wales 3: The population status of birds in wales. Birds in Wales, 13, 3–31.

Kelling, S., Johnston, A., Fink, D., Ruiz-Gutierrez, V., Bonney, R., Bonn, A., Fernandez, M., Hochachka, W., Julliard, R., Kraemer, R. & others. (2018) Finding the signal in the noise of citizen science observations. bioRxiv, 326314.

Kellner, K. (2018) jagsUI: A Wrapper Around ‘Rjags’ to Streamline ‘JAGS’ Analyses.

Kéry, M., Royle, J.A., Schmid, H., Schaub, M., Volet, B., Haefliger, G. & Zbinden, N. (2010) Site-occupancy distribution modeling to correct population-trend estimates derived from opportunistic observations. Conservation Biology, 24, 1388–1397.

Kirsop-Taylor, N. (2019) The means, motive and opportunity of devolved policy responses to an ecosystem approach. British Politics, 1–20.

Lawton, J.H. (1993) Range, population abundance and conservation. Trends in Ecology and Evolution, 8, 409–413.

Lomba, A., Guerra, C., Alonso, J., Honrado, J.P., Jongman, R. & McCracken, D. (2014) Mapping and monitoring high nature value farmlands: Challenges in european landscapes. Journal of Environmental Management, 143, 140–150.

Luebcke, W. (1977) 17 jahre beringung an einem schlafplatz der grauammer (emberiza calandra). Vogelkundliche Hefte Edertal, 17, 57–72.

Martay, B., Pearce-Higgins, J.W., Harris, S.J. & Gillings, S. (2018) Monitoring landscape-scale environmental changes with citizen scientists: Twenty years of land use change in great britain. Journal for nature conservation, 44, 33–42.

Meyer, C., Jetz, W., Guralnick, R.P., Fritz, S.A. & Kreft, H. (2016) Range geometry and socio-economics dominate species-level biases in occurrence information. Global Ecology and Biogeography, 25, 1181–1193.

Meyer, C., Kreft, H., Guralnick, R. & Jetz, W. (2015) Global priorities for an effective information basis of biodiversity distributions. Nature Communications, 6, 8221.

Morrison, C.A., Robinson, R.A., Clark, J.A., Risely, K. & Gill, J.A. (2013) Recent population declines in afro-palaearctic migratory birds: The influence of breeding and non-breeding seasons. Diversity and Distributions, 19, 1051–1058.

Natural England. (2014) National Character Area Profiles.

Newson, S.E., Massimino, D., Johnston, A., Baillie, S.R. & Pearce-Higgins, J.W. (2013) Should we account for detectability in population trends? Bird Study, 60, 384–390.

Newson, S.E., Moran, N.J., Musgrove, A.J., Pearce-Higgins, J.W., Gillings, S., Atkinson, P.W., Miller, R., Grantham, M.J. & Baillie, S.R. (2016) Long-term changes in the migration phenology of UK breeding birds detected by large-scale citizen science recording schemes. Ibis, 158, 481– 495.

Outhwaite, C.L., Gregory, R.D., Chandler, R.E., Collen, B. & Isaac, N.J. (2020) Complex long-term biodiversity change among invertebrates, bryophytes and lichens. Nature Ecology & Evolution, 4, 384–392.

Pearce-Higgins, J.W., Ockendon, N., Baker, D.J., Carr, J., White, E.C., Almond, R.E., Amano, T., Bertram, E., Bradbury, R.B., Bradley, C. & others. (2015) Geographical variation in species’ population responses to changes in temperature and precipitation. Proceedings of the Royal Society B, 282, 20151561.

Perkins, A.J., Maggs, H.E., Watson, A. & Wilson, J.D. (2011) Adaptive management and targeting of agri-environment schemes does benefit biodiversity: A case study of the corn bunting emberiza calandra. Journal of Applied Ecology, 48, 514–522.

Plummer, M. (2003) JAGS: A program for analysis of Bayesian graphical models using Gibbs sampling. Proceedings of the 3rd international workshop on distributed statistical computing Vienna, Austria.

R Core Team. (2018) R: A Language and Environment for Statistical Computing. R Foundation for Statistical Computing, Vienna, Austria.

Roberts, R., Donald, P. & Green, R. (2007) Using simple species lists to monitor trends in animal populations: New methods and a comparison with independent data. Animal Conservation, 10, 332–339.

Robinson, R.A. (2010) State of bird populations in britain and ireland. Silent summer: The state of wildlife in britain and ireland (ed N. Maclean), pp. 281–318. Cambridge University Press.

Robinson, O.J., Ruiz-Gutierrez, V. & Fink, D. (2018) Correcting for bias in distribution modelling for rare species using citizen science data. Diversity and Distributions, 24, 460–472.

Robinson, O.J., Ruiz-Gutierrez, V., Reynolds, M.D., Golet, G.H., Strimas-Mackey, M. & Fink, D. (2020) Integrating citizen science data with expert surveys increases accuracy and spatial extent of species distribution models. Diversity and Distributions.

Rose-Ackerman, S. (1994) Environmental policy and federal structure: A comparison of the united states and germany. Vande. L. Rev., 47, 1587.

Roy, H.E., Adriaens, T., Isaac, N.J., Kenis, M., Onkelinx, T., Martin, G.S., Brown, P.M., Hautier, L., Poland, R., Roy, D.B. & others. (2012) Invasive alien predator causes rapid declines of native european ladybirds. Diversity and Distributions, 18, 717–725.

Roy, D., Harding, P., Preston, C. & Roy, H. (2014) Celebrating 50 Years of the Biological Records Centre. NERC/Centre for Ecology & Hydrology.

Schmeller, D.S., Henry, P.-Y., Julliard, R., Gruber, B., Clobert, J., Dziock, F., Lengyel, S., Nowicki, P., Deri, E. & Budrys, E. (2009) Advantages of volunteer-based biodiversity monitoring in europe. Conservation Biology, 23, 307–316.

Sorte, F.A.L. & Somveille, M. (2020) Survey completeness of a global citizen-science database of bird occurrence. Ecography, 43, 34–43.

Specht, H. & Lewandowski, E. (2018) Biased assumptions and oversimplifications in evaluations of citizen science data quality. Bulletin of the Ecological Society of America, 99, 251–256.

Stanbury, A., Davies, M., Grice, P., Gregory, R. & Wotton, S. (2010) The status of the cirl bunting in the uk in 2009. British Birds, 103, 702.

Stevens, M., Murn, C. & Hennessey, R. (2019) Population change of common buzzards buteo buteo in central southern england between 2011 and 2016. Bird Study, 66, 378–389.

Stevens, M., Murn, C. & Hennessey, R. (2020) Population change of red kites milvus milvus in central southern england between 2011 and 2016 derived from line transect surveys and multiple covariate distance sampling. Acta Ornithologica, 54, 243–254.

van Strien, A.J., Pannekoek, J. & Gibbons, D.W. (2001) Indexing european bird population trends using results of national monitoring schemes: A trial of a new method. Bird Study, 48, 200–213.

van Strien, A.J., van Swaay, C.A. & Termaat, T. (2013) Opportunistic citizen science data of animal species produce reliable estimates of distribution trends if analysed with occupancy models. Journal of Applied Ecology, 50, 1450–1458.

van Swaay, C.A., Dennis, E., Schmucki, R., Sevilleja, C., Balalaikins, M., Botham, M., Bourn, N., Brereton, T., Cancela, J., Carlisle, B. & others. (2019) The Eu Butterfly Indicator for Grassland Species: 1990-2017. Butterfly Conservation Europe & ABLE/eBMS.

van Swaay, C.A., Nowicki, P., Settele, J. & van Strien, A.J. (2008) Butterfly monitoring in europe: Methods, applications and perspectives. Biodiversity and Conservation, 17, 3455–3469.

Walker, J. & Taylor, P. (2017) Using eBird data to model population change of migratory bird species. Avian Conservation and Ecology, 12.

Welsh Government. (2017) Natural Resources Policy. Welsh Government.

Wotton, S.R., Bladwell, S., Mattingley, W., Morris, N.G., Raw, D., Ruddock, M., Stevenson, A. & Eaton, M.A. (2018) Status of the hen harrier circus cyaneus in the uk and isle of man in 2016. Bird Study, 65, 145–160.

Zipkin, E.F., Rossman, S., Yackulic, C.B., Wiens, J.D., Thorson, J.T., Davis, R.J. & Grant, E.H.C. (2017) Integrating count and detection–nondetection data to model population dynamics. Ecology, 98, 1640–1650.

